# HDAC4/5 regulate epidermal barrier function by modulating the epigenetic landscape of human keratinocytes

**DOI:** 10.64898/2026.08.13.741670

**Authors:** Chloé Nguyen Van, Stéphanie Denis, Sébastien Cadau, Nicolas Pelletier, Valérie André, Jérôme Lamartine

## Abstract

Keratinocyte proliferation and differentiation are essential to produce the stratified structure of the epidermis and maintain its barrier function. These processes are regulated by complex mechanisms including epigenetic regulation. In this study, we evaluated the role of HDAC4/5, two class IIa histone deacetylases, in the epigenetic regulation of proliferative and differentiated human keratinocytes using dedicated 2D and 3D *in vitro* models. Our findings demonstrate that chemical inhibition or shRNA-mediated knock-down of HDAC4 impair keratinocyte proliferation notably through increased H3K27 acetylation and subsequent transcriptional activation of the cell cycle inhibitor gene *BTG2*. Interestingly, HDAC4/5 inhibition alters H3K27 acetylation landscape in proliferating keratinocytes, whereas the epigenetic identity of differentiated keratinocytes is much less affected. Inhibiting HDAC4/5 in 3D epidermis models resulted in reduced epidermal thickness and impaired barrier function linked to alteration in the lipid composition of the *stratum corneum*. Furthermore, analysis of several well-established skin aging markers revealed that reconstructed human epidermis treated with the HDAC4/5 inhibitor exhibit molecular and functional characteristics consistent with an aged-epidermis. Collectively, our results demonstrate that HDAC4/5 are essential for maintaining epidermal homeostasis and pave the way for the development of innovative models of skin aging based on the modulation of histone acetylation.

## INTRODUCTION

The epidermis plays a major role in maintaining barrier integrity notably through its highly renewable capacity. This process relies on a dynamic balance between keratinocyte proliferation and differentiation, which contributes to preserving tissue architecture and thickness (Al-Dhubaibi et al. 2025). Moreover, differentiated keratinocytes display a specific metabolic program, mainly through their ability to synthesize and organize lipids within the *stratum corneum*. These lipids form a highly organized multilamellar matrix surrounding corneocytes and thus contribute to the epidermal barrier function (Van Smeden et al. 2014).

The switch from a proliferative to a differentiated cell state required to maintain epidermal homeostasis, relies on fine-tuned gene regulation in which epigenetic mechanisms play a fundamental role by adjusting chromatin compaction and consequently gene accessibility (Botchkarev et al. 2012). Among the epigenetic marks, histone acetylation and deacetylation, respectively catalyzed by histone acetyltransferases (HATs) and histone deacetylases (HDACs), are known to regulate epidermis development and functions (Kang et al. 2019). The HDAC family comprises two subfamilies: the zinc dependent HDACs composed of 11 members and the NAD+ dependent HDACs composed of 7 sirtuins (Finkel et al. 2009; Haberland et al. 2009). These enzymes catalyze the elimination of acetyl groups from the histone lysine residues resulting in an increased positive charge of histones, which condense into heterochromatin, thereby decreasing gene transcription (Bannister and Kouzarides 2011). A study has demonstrated the transcriptional expression of all HDACs in keratinocytes (Sanford et al. 2016). Among them, HDAC2 and HDAC9 were preferentially expressed in basal keratinocytes, whereas most HDACs transcripts were more abundant in differentiated keratinocytes, with HDAC3 showing the highest expression levels in both populations. Regarding the physiological roles of HDACs in the epidermis, their regulation of cell cycle genes has been well described and may explain the growing interest of these enzymes in the contexts of cancer, wound healing and tissue repair processes (Saunders et al. 1999; Spallotta et al. 2013).

Despite these findings, the contribution of individual HDAC family members to epidermal homeostasis remains poorly characterized, with most studies focusing on HDAC1 and HDAC2. Deletion of both HDAC1 and HDAC2 in mice embryos impairs epidermal development (LeBoeuf et al. 2010), whereas co-deletion of HDAC1 and HDAC2 in adult mice leads to reduced proliferation, impaired differentiation and increased apoptosis (Zhu et al. 2022). Although HDAC1/2 are now recognized as regulators of epidermal development, the roles of other HDAC family members remain largely unexplored as most studies in human keratinocytes have relied on pan-HDAC inhibitors (Markova et al. 2007). Among the less characterized HDACs, emerging evidence suggests a role for HDAC4/5 in skin biology. HDAC5 has recently been implicated in skin homeostasis during wound healing (Zhang et al. 2025), yet limited knowledge is available regarding the function of HDAC4 in the skin. The only reported findings showed HDAC4 protective role in the context of cell senescence in human dermal fibroblasts (Di Giorgio et al. 2021; Lee et al. 2022) and its decreased activity during skin aging (Lee et al. 2021).

Therefore, the precise roles of HDAC4 and HDAC5 in human epidermal keratinocytes and specifically how HDAC4/5 modulate the epigenetic landscape to maintain epidermal homeostasis need to be elucidated. To address this question, we focused our study on HDAC4/5 function in human epidermal homeostasis by using chemical inhibition of HDAC4/5 and RNA-interference mediated knock-down of HDAC4 in complementary 2D and 3D keratinocyte models. We showed that HDAC4/5 inhibition alters epidermal homeostasis by impairing cell proliferation and inducing abnormal keratinocyte differentiation, as well as compromising skin barrier function mainly due to alterations in lipids synthesis. We further demonstrated that HDAC4/5 inhibition significantly alters the histone H3 lysine 27 acetylation (H3K27ac) landscape in proliferating keratinocytes, whereas the epigenetic identity of differentiated keratinocytes is much less affected. Together, this study provides new insights into the role of histone acetylation in regulating keratinocyte differentiation and, more broadly, in maintaining the vital epidermal barrier function.

## RESULTS

### HDAC4/5 inhibition alters cell proliferation in monolayer keratinocytes and reconstructed human epidermis

To investigate the role of HDAC4/5 in human keratinocytes, we treated immortalized keratinocytes (N/TERT1) with LMK235, a selective HDAC4/5 inhibitor with a high efficiency on HDAC4 activity (Li et al. 2018; Tang et al. 2024). To evaluate its efficiency on HDAC activity, we first assessed the acH3 protein levels in monolayer N/TERT1 cultures and found a significant increase of acH3 after treatment with 1 µM LMK235 compared with untreated cells (Figure 1a and 1b). We also observed that LMK235 reduced global HDAC activity by 35% (Figure 1c). These results show the effectiveness of LMK235 in inhibiting HDAC activity and consequently, increasing acH3 levels.

**Figure 1.**
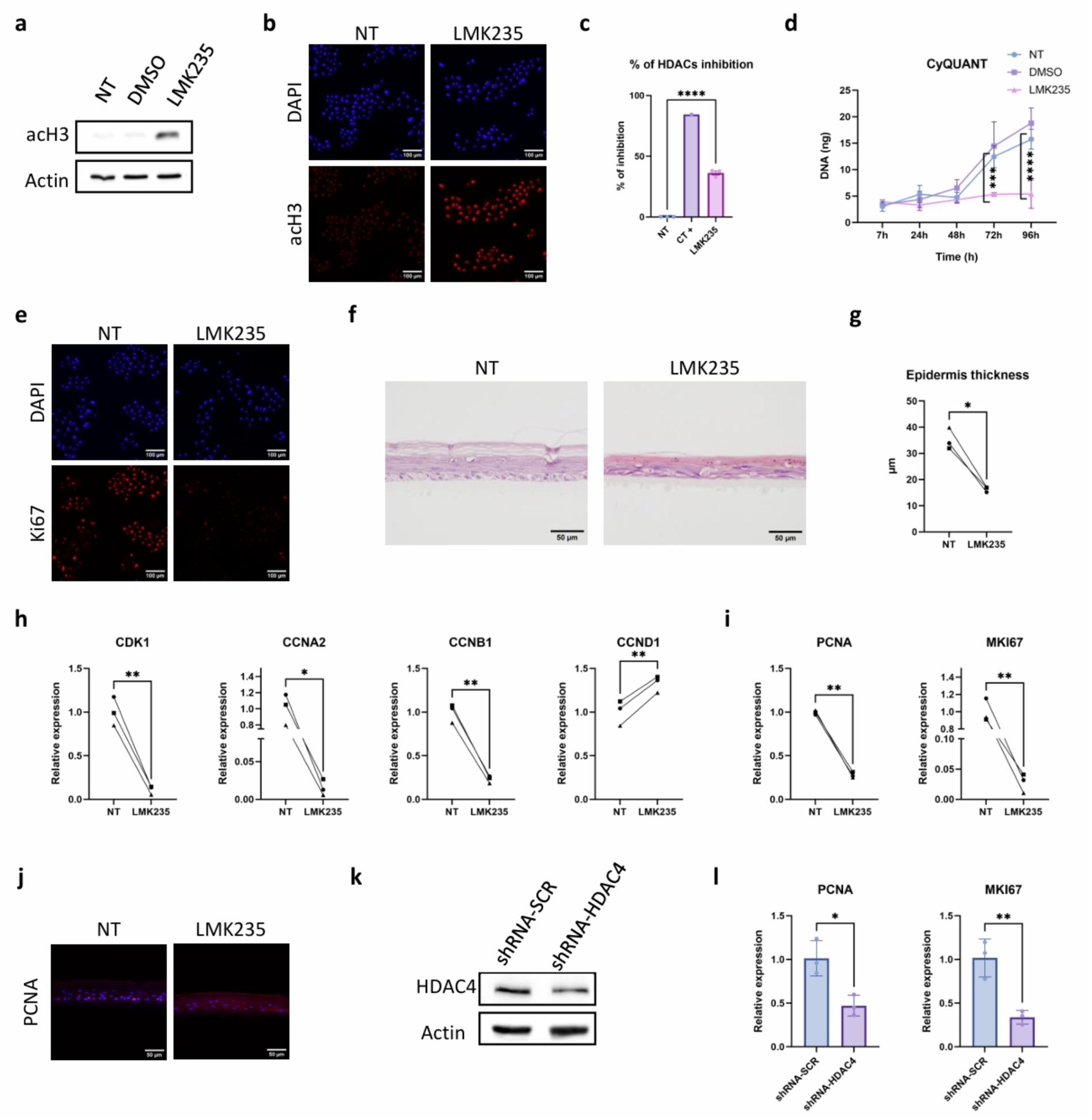
HDAC4/5 inhibition increases acH3 levels and impairs keratinocyte proliferation in 2D and 3D models. (a) Western-blot analysis of acH3 in N/TERT1 monolayer cells treated or not with either 1 µM LMK235 or DMSO for 24h. Actin was used as a loading control. (b) Representative immunofluorescence images of acH3 in N/TERT1 monolayer cells treated or not with 1 µM LMK235 for 24h. DAPI was used to stain nuclei. Bars=100 µM. (c) Percentage of HDACs inhibition determined by HDACs activity measurement in N/TERT1 monolayer cells treated or not with 1 µM LMK235. TSA pan-HDAC inhibitor is used as a positive control of total HDACs inhibition (**** P<0.0001, n=3, Student’s t-test). (d) Proliferation profile of N/TERT1 monolayer cells treated or not with either 1 µM LMK235 or DMSO. DNA concentration was measured 7h, 24h, 48h, 72h and 96h after the beginning of the treatment (*** P<0.001, **** P<0.0001, n=3, ANOVA2). (e) Representative immunofluorescence images of the proliferation marker Ki67 in N/TERT1 monolayer cells treated or not with 1 µM LMK235 for 24h. DAPI was used to stain nuclei. Bars=100 µM. (f) Representative H&E images of reconstructed human epidermis treated or not with 1 µM LMK235. Bars= 50 µM. (g) Quantification of epidermis thickness in reconstructed human epidermis treated or not with 1 µM LMK235 (* P<0.05, N=3, Student’s t-test). (h) Relative mRNA expression for *CDK1*, *CCNA2*, *CCNB1* and *CCND1* in reconstructed human epidermis treated or not with 1 µM LMK235. Data were normalized to housekeeping genes and expressed relative to the untreated condition (* P<0.05, ** P<0.01, N=3, Student’s t-test). (i) Relative mRNA expression for *PCNA* and *MKI67* in reconstructed human epidermis treated or not with 1 µM LMK235. Data were normalized to housekeeping genes and expressed relative to the untreated condition (** P<0.01, N=3, Student’s t-test). (j) Representative immunofluorescence images of the proliferation marker PCNA in reconstructed human epidermis treated or not with 1 µM LMK235. DAPI was used to stain nuclei. Bars=50 µM. (k) Western-blot analysis of HDAC4 in N/TERT1 monolayer cells transduced with shRNA-Scrambled (shRNA-SCR, control) or shRNA-HDAC4. Actin was used as a loading control. (l) Relative mRNA expression for *PCNA* and *MKI67* in N/TERT1 monolayer cells transduced with shRNA-SCR or shRNA-HDAC4. Data were normalized to housekeeping genes and expressed relative to the shRNA-Scrambled condition (* P<0.05, ** P<0.01, n=3, Student’s t-test).

We then investigated the impact of HDAC4/5 inhibition on keratinocyte proliferation, by measuring total DNA concentration. While DNA concentration increased over time in untreated cells, it remained unchanged in HDAC4/5-inhibited cells, suggesting that the cell cycle was strongly impacted (Figure 1d). These results were confirmed by a decrease of Ki67 levels, a proliferative marker (Figure 1e) (Supplementary Figure S1a). We next explored whether this alteration of proliferation after LMK235 treatment was also effective in 3D models. We generated reconstructed human epidermis (RHE) using human primary keratinocytes (HPK) treated or not with LMK235. We observed that HDAC4/5 inhibition impaired epidermal development and led to an abnormal basal layer (Figure 1f). At the morphological level, basal cells appeared to lose their characteristic cuboidal shape and adopt a more elongated morphology, similar to that observed in cells of the suprabasal layers. In addition, the epidermal thickness was significantly reduced by approximately 50%, likely reflecting impaired cell renewal caused by altered cell proliferation (Figure 1g). To confirm the alteration of cell proliferation, the transcript levels of several cyclins and proliferation markers were analyzed by RT-qPCR. The LMK235 treatment decreased the expression of genes encoding CDK1, cyclin A2 (*CCNA2)* and cyclin B1 (*CCNB1)*, whereas the expression of the gene encoding cyclin D1 (*CCND1)* increased suggesting cell-cycle arrest and a cell accumulation in the G1 phase (Figure 1h). The antiproliferative effect of LMK235 in RHE was further supported by the reduced expression of the genes encoding PCNA and Ki67 (*MKI67*) (Figure 1i) together with a lower number of PCNA-positive cells detected by immunostaining (Figure 1j). To clarify the specific and direct role of HDAC4 on keratinocyte proliferation, a shRNA-mediated knock-down of HDAC4 cells was performed in N/TERT1 cells (Figure 1k). Similar to the pharmacological inhibition, silencing of HDAC4 led to a significant decrease in *PCNA*, *MKI67*, *CDK1*, *CCNA2* and *CCNB1* transcripts levels (Figure 1l) (Supplementary Figure S1b).

### HDAC4/5 inhibition reshapes the transcriptional profile and impairs differentiation in reconstructed human epidermis

To obtain a global and unbiased understanding of the impact of HDAC4/5 inhibition, we performed a transcriptomic analysis by RNA-seq on RHE treated or not with LMK235. Based on the sample-to-sample Euclidean distance, a strong transcriptomic signature of HDAC inhibition was identified (Figure 2a), consistent with the established role of these enzymes as epigenetic regulators and reflecting the broad impact of modulating chromatin-associated processes. A total of 3 614 Differentially Expressed Genes (DEGs) was identified in the human epidermal equivalents following LMK235 treatment with a similar proportion of up-regulated genes (1766 genes) and down-regulated genes (1848 genes) (|log2FoldChange|>1, padj<0.05) (Figure 2b). The functional annotation of these DEGs was performed and the top 8 annotations associated with down-regulated and up-regulated genes are presented in Figure 2c and Figure 2d, respectively. Interestingly, HDAC4/5 inhibition decreased the expression of genes involved in pathways related to cell proliferation corroborating our previous observations.

**Figure 2.**
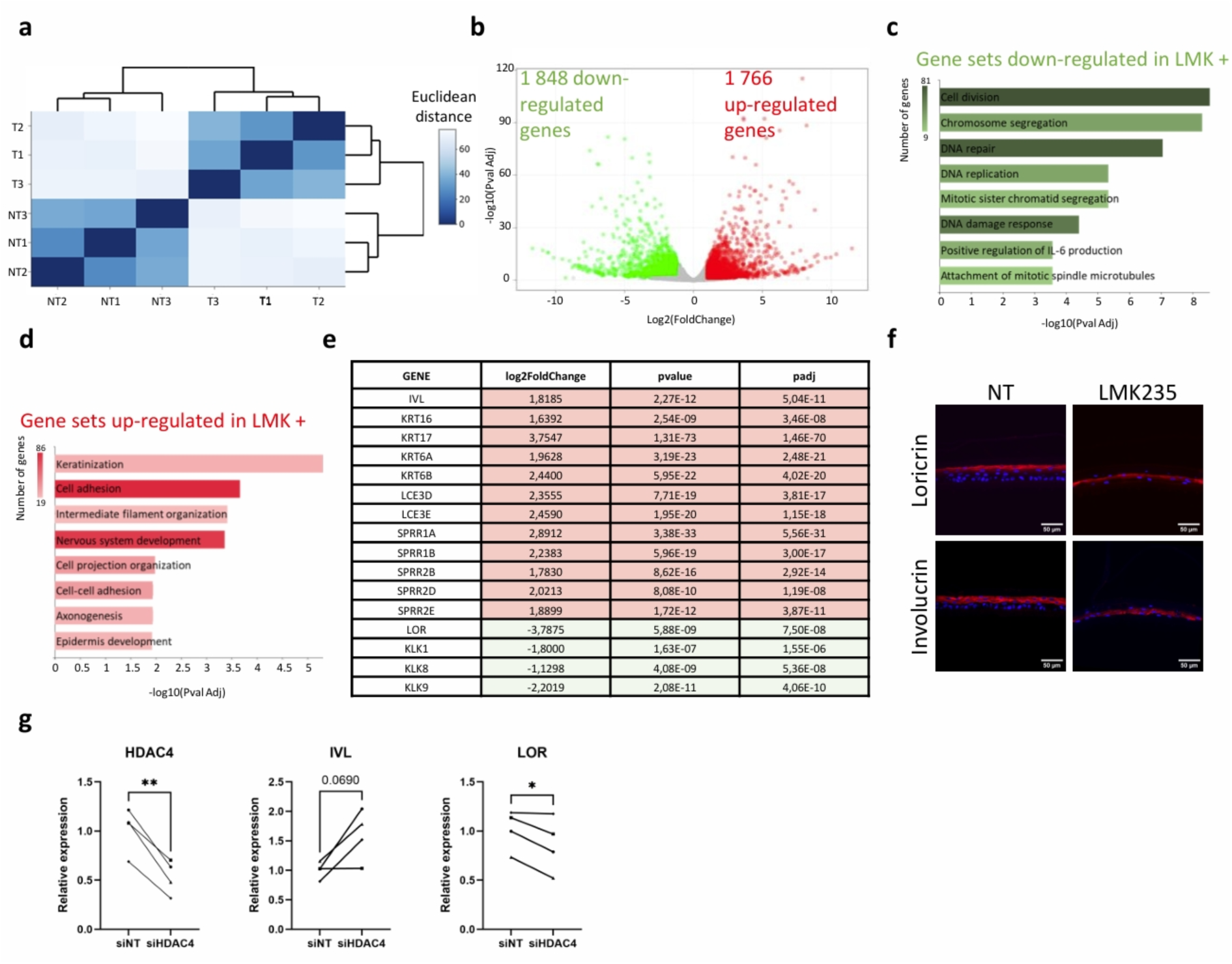
HDAC4/5 inhibition reshapes transcriptional profiles and enhances keratinization leading to accelerated but abnormal and incomplete differentiation in RHE. (a) Heat map of the sample-to-sample Euclidean distance of RNA-seq profiles in reconstructed human epidermis treated or not with 1 µM LMK235. Color scale represents the Euclidean distance (N=3). (b) Volcano plot of the transcriptional profiles of the reconstructed human epidermis treated or not with 1 µM LMK235. The 1848 green and 1766 red dots represent genes whose expressions were respectively down-regulated and up-regulated in the 1 µM LMK235 treated group (N=3) (c) Bar chart showing the top 8 gene annotations of down-regulated genes in the 1 µM LMK235 treated group (N=3) (d) Bar chart showing the top 8 gene annotations of up-regulated genes in the 1 µM LMK235 treated group (N=3) (e) Table showing the differentially expressed genes involved in the keratinization pathway in reconstructed human epidermis treated or not with 1 µM LMK235. The table lists gene names, log2FoldChange, p values and adjusted p values (padj). (f) Representative immunofluorescence images of two markers of keratinocytes differentiation, loricrin and involucrin in reconstructed human epidermis treated or not with 1 µM LMK235. DAPI was used to stain nuclei. Bars=50 µM. (g) Relative mRNA expression for *HDAC4*, *IVL* and *LOR* in reconstructed human epidermis generated from HPK transfected with non-targeting siRNA (siNT) and HDAC4-targeting siRNA (siHDAC4). Data were normalized to housekeeping genes and expressed relative to the siNT condition (* P<0.05, ** P<0.01, N=4, Student’s t-test).

Conversely, LMK235 treatment increased the expression of genes involved in epidermal development and keratinization. Analysis of the up-regulated genes associated with these biological pathways revealed increased expression of involucrin (IVL), several keratin genes (*KRT6A*, *KRT6B*, *KRT6C*, *KRT16* and *KRT17*), late cornified envelop genes (*LCE3D* and *LCE3E*) and *SPRR* genes (Figure 2e). Interestingly these genes are known to be up-regulated in response to skin injury and impaired barrier function (Bergboer et al. 2011; De Cid et al. 2009; De Koning et al. 2012; Zhang et al. 2019). Moreover, SPRR and LCE proteins are structurally related to loricrin and have been shown to be up-regulated in loricrin knock-out models, where they act as alternative protein to compensate for altered epidermal proprieties including differentiation processes required for epidermal barrier integrity (Ishitsuka and Roop 2020). Notably, significant down-regulation of the terminal differentiation markers loricrin (*LOR*) as well as kallikreins (*KLKs*) was observed in RHE following LMK235 treatment (Figure 2e). Furthermore, we observed an abnormal localization of involucrin and loricrin with a middle layer expression for both proteins (Figure 2f). Taken together, these results suggest a complex effect of HDAC4/5 inhibition leading to a disorganized differentiation process that may either contribute to or result from an altered epidermal barrier function. This hypothesis is strengthened by the reduced number of keratohyalin granules in the uppermost *stratum granulosum* (Supplementary Figure S1i) of treated RHE, which is considered to be an adaptative response to barrier disruption (Hirabayashi et al. 2017).

To specify the relative contribution of HDAC4 inhibition in the phenotype of LMK235-treated RHE, we generated RHE using HPK transfected with non-targeting siRNA (siNT) and HDAC4-targeting siRNA (siHDAC4). After 6 days of culture at air-liquid interface, we obtained a thinner epidermis for siHDAC4 compared to siNT RHE (Supplementary Figure S1c and S1d), which confirms the phenotype observed with LMK235. RT-qPCR analysis of these tissues confirmed a 50 % reduction of the *HDAC4* transcript levels (P<0.01), a 60 % induction of the *IVL* gene (yet not statistically significant) and a 15 % repression of the *LOR* gene (P<0.05) (Figure 2g), similar to what was observed in RHE treated with LMK235.

### HDAC4/5 inhibition alters epidermal barrier function and impairs lipids synthesis

Since the transcriptomic data suggest that the epidermal differentiation is affected by HDAC4/5 inhibition, we evaluated the barrier integrity of treated reconstructed human epidermis. LMK235 significantly increased trans-epidermal water loss (TEWL) and decreased trans-epidermal electrical resistance (TEER) (Figure 3a) suggesting an impairment of the epidermal barrier function after HDAC4/5 inhibition. To investigate whether this alteration is associated with abnormal *stratum corneum* lipid composition, we generated a full-thickness skin model (RHS) to provide a more physiologically relevant model with a complete terminal differentiation and a fully developed *stratum corneum*. As for RHE, HDAC4/5 inhibition led to a decrease of epidermal thickness (Figure 3b). We then performed lipidomic analysis by mass-spectrometry on the RHS. The analysis of the lipid composition revealed a significant decrease in ceramides NS (non-hydroxy fatty acid sphingosine), AS (α-hydroxy fatty acid sphingosine) and EOS (esterified ω-hydroxy fatty acid sphingosine) upon HDAC4/5 inhibition (Figure 3c). As the *stratum corneum* lipid matrix is essential to maintain the epidermal barrier integrity (Schild et al. 2024), these results may explain the observed impairment of water permeability. To further investigate the alteration of lipid expression, we analyzed the transcript levels of key enzymes involved in the ceramide synthesis pathway. We observed a significant decrease in *ELOVL4* transcripts encoding a protein involved in the elongation of very long fatty acid chain, as well as in *CERS2* and *CERS3* transcripts, which encode two ceramide synthases (Figure 3d). Interestingly, *ASAH1* mRNA levels, encoding a ceramidase, were decreased following LMK235 treatment. This reduction could reflect a compensatory mechanism aimed at preserving the remaining lipid stores. To determine whether HDAC4/5 inhibition affects the ultra-structure of the *stratum corneum*, we performed transmission electronic microscopy on RHE treated or not with LMK235. We observed a decrease in the number of lipid lamellae, which also appeared thinner upon HDAC4/5 inhibition (Figure 3e and Supplementary Figure S1e). This was associated with an overall compaction of the *stratum corneum*. Nevertheless, in both conditions, lipid droplets, cholesterol crystals and lamellar bodies remained visible. Overall, these data indicate that HDAC4/5 inhibition leads to an alteration of the barrier integrity associated with an impairment in lipid biosynthesis pathway and altered structure of the *stratum corneum*.

**Figure 3.**
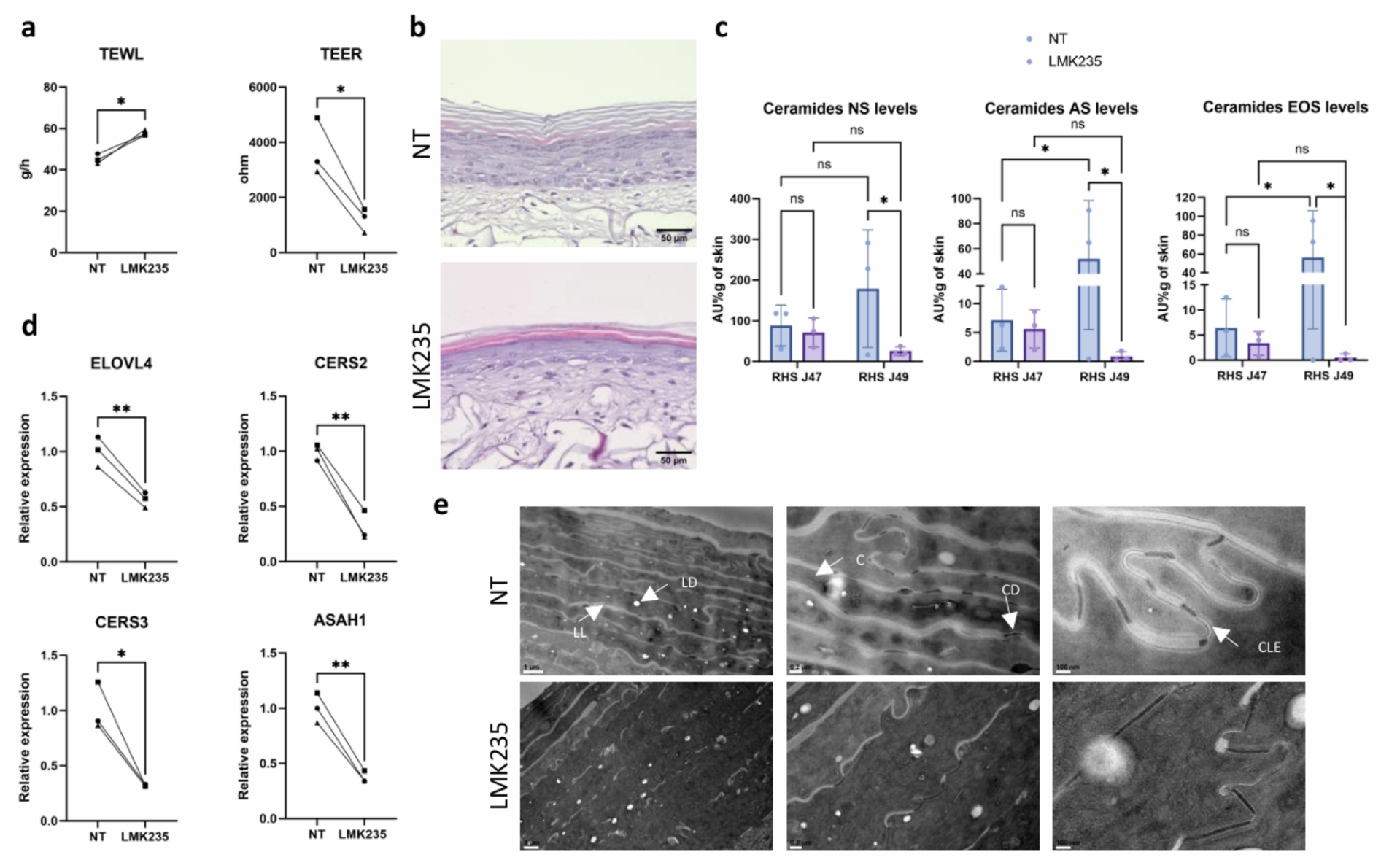
HDAC4/5 inhibition impairs barrier function through altered lipid synthesis and lipid lamellae architecture. (a) Measurement of trans-epidermal water loss (TEWL) and trans-epidermal electrical resistance (TEER) in reconstructed human epidermis treated or not with 1 µM LMK235. For each experimental condition and each biological replicate, measurements were performed in triplicate, and the mean value was calculated. (* P<0.05, N=3, Student’s t-test). (b) Representative H&E images of full thickness reconstructed human skin treated or not with 1 µM LMK235. Bars= 50 µM. (c) NS, AS and EOS ceramides levels in full thickness reconstructed human skin after 47 and 49 days of culture, treated or not with 1 µM LMK235. (* P<0.05, n=3, ANOVA2). (d) Relative mRNA expression for *ELOVL4*, *CERS2*, *CERS3* and *ASAH1* in reconstructed human epidermis treated or not with 1 µM LMK235. Data were normalized to housekeeping genes and expressed relative to the untreated condition (* P<0.05, ** P<0.01, N=3, Student’s t-test). (e) Transmission Electron Microscopy (TEM) representative images of the stratum corneum of reconstructed human epidermis treated or not with 1 µM LMK235. LL: lipid lamellae, LD: lipid droplets, CD: corneodesmosomes, C: corneocytes, CLE: corneocyte lipid envelope (N=1). Bars= 1 µM, 0,2 µM and 100 nm (from left to right column).

### Proliferative and differentiated keratinocytes exhibit distinct H3K27ac epigenetic landscapes

To better understand how histone acetylation contributes to the proliferation/differentiation balance of keratinocytes during epidermal development, we performed H3K27ac ChIP-seq analysis in proliferative or differentiated N/TERT1 treated or not with LMK235. Interestingly, most of the 250 peaks with the highest variance across experimental conditions displayed higher H3K27ac levels in proliferative non-treated control cells compared to differentiated untreated cells (Figure 4a). An increase of H3K27ac following LMK235 treatment was observed for most of these peaks although HDAC4/5 inhibition did not disrupt the global epigenetic identity of the two cellular states. Analysis of the differentially bound peaks between differentiated and proliferative treated cells revealed that LMK235 treatment preferentially increased H3K27ac in proliferative cells compared to differentiated cells (Figure 4b). Indeed, whereas 40,114 differentially bound peaks mapping to 15,001 genes were identified in proliferative cells upon LMK235 treatment, only 15,665 differentially bound peaks mapping to 7,545 genes were observed in differentiated cells (padj<0.01).

**Figure 4.**
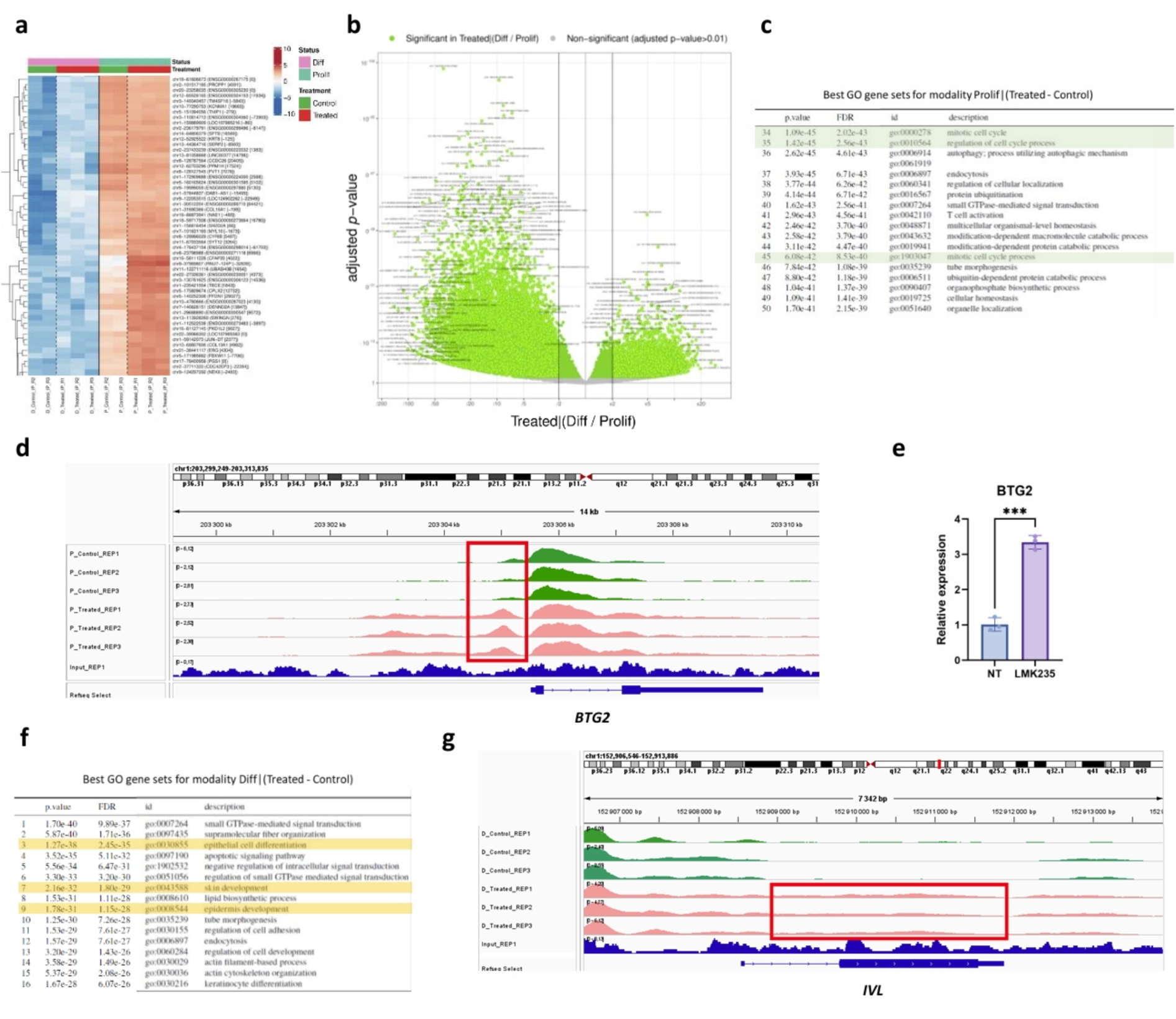
Distinct epigenetic landscapes underlie proliferative and differentiated states that are modulated by HDAC4/5 activity without a complete disruption of cell identity. (a) Heatmap of several of the 250 peaks with highest variance in the H3K27ac ChIP-seq experiment (format: “genomic location (closest gene [distance])”, with a positive distance reflecting a peak downstream of the transcription start site). Color represents the intensity of the centered (but unscaled) peak signal that goes, for each peak, from low (blue), to medium (white) to high (ref). For each of the top subclusters, at most 3 GO terms that, combined, best overlap with the genes closest to the corresponding peaks, are displayed. (b) Volcano-plot of differentially bound peaks between differentiated treated cells and proliferative treated cells (Treated|(Diff/Prolif)). Annotation format: “genomic location(closest gene[distance])” with a positive distance reflecting a peak downstream of the transcription start site. The vertical black lines delimit the 2-fold change effects. (c) Table listing the top 34-50 most significant Gene Ontology gene sets identified using the GREAT method for the proliferative treated cells compared with the proliferative control cells. (d) Integrative Genomics Viewer (IGV) visualization of H3K27ac ChIP-seq signal at the *BTG2* locus in proliferative treated cells (pink) or proliferative control cells (green) (n=3). (e) Relative mRNA expression for *BTG2* in N/TERT1 monolayer cells treated or not with 1 µM LMK235 for 24h. Data were normalized to housekeeping genes and expressed relative to the untreated condition (*** P<0.001, n=3. Student’s t-test). (f) Table listing the top 1-16 most significant Gene Ontology gene sets identified using the GREAT method for the differentiated treated cells compared with the differentiated control cells. (g) Integrative Genomics Viewer (IGV) visualization of H3K27ac ChIP-seq signal at the *IVL* locus in differentiated treated cells (pink) or differentiated control cells (green) (n=3).

Overall, these results suggest that the proliferative cells contain more accessible genomic regions and exhibit greater epigenetic plasticity. This permissive chromatin state is possibly important to ensure cell fate flexibility, in comparison to differentiated keratinocytes, which have already committed to a specific cellular program, thereby displaying less accessible genomic region. Moreover, the limited effect of LMK235 treatment on differentiated cells may result from the fact that the genomic regions required to sustain this cellular state are already highly acetylated and therefore saturated with acetyl groups.

### Keratinocyte proliferation and differentiation are epigenetically regulated

Gene ontology (GO) analysis of the gene sets associated with the differentially bound peaks between the untreated and treated proliferative cells revealed several pathways related to cell division (Figure 4c). As LMK235 treatment altered proliferation, we investigated whether some cell cycle inhibitors could be directly targeted by HDAC4/5. We identified *BTG2*, a tumor suppressor gene encoding an antiproliferative protein (Yuniati et al. 2019), as one of these potential targets. Indeed, we observed a broad H3K27ac-enriched domain upstream of the transcription start site (TSS), overlapping with an enhancer region. This suggests that HDAC4/5 inhibition increases the activity of this regulatory region, potentially contributing to increased gene transcription (Figure 4d). Consistently, transcript analysis showed an increase in *BTG2* mRNA in N/TERT1 cells treated with LMK235 (Figure 4e) and N/TERT1 cells transduced with shRNA-HDAC4 (Supplementary Figure S1f). Taken together, these results suggest that *BTG2* may be a direct target of HDAC4 and possibly HDAC5 and that under HDAC4/5 inhibition conditions, increased histone acetylation at the *BTG2* locus promotes cell cycle arrest. The downregulation of cyclin transcripts previously observed may thus be a consequence of *BTG2* increased transcription (Figure 1h). Interestingly, the downregulation of cell-cycle genes in LMK235 treated cells, could also potentially result from hypoacetylated regions located at the transcription start sites, as observed for *MKI67* through the H3K27ac ChIP-seq analysis (Supplementary Figure S1h).

Gene ontology of the gene set associated with the differentially bound peaks between the untreated and treated differentiated cells pointed to several pathways related to epidermis development (Figure 4f). As we previously observed an increase in *IVL* transcript levels after treatment, we investigated whether *IVL* could also be directly targeted by HDAC4/5. Visualization of H3K27ac ChIP-seq signal revealed a broad enrichment across the *IVL* gene body (Figure 4g). This finding suggests that HDAC4/5 inhibition leads to multiple H3K27ac marked nucleosomes, which have been shown to occur at cis-regulatory elements, along gene bodies and in intragenic regions, resulting in extensive domain of active chromatin (Di Giorgio et al. 2021; Paldi et al. 2026). The absence of H3K27ac peaks near to the initiation start site suggests that the promoter was already active under the untreated condition, due to the differentiated state of the cells. Collectively, these results show that keratinocyte proliferation and differentiation are epigenetically regulated and that cell cycle inhibitors as well as differentiation markers such as involucrin are likely to be directly regulated by HDAC4/5.

### HDAC4 is downregulated during aging and HDAC4 inhibition leads to an aging-phenotype

We have demonstrated that HDAC4/5 inhibition impairs cell proliferation, decreases epidermal thickness and alters barrier function mainly through reduced lipid synthesis in the *stratum corneum*. Interestingly, all of these events are hallmarks of epidermal aging. We therefore wondered whether HDAC4 and HDAC5 levels could be affected during intrinsic aging. Interestingly, we observed decreased HDAC4 proteins and transcripts levels in human primary keratinocytes extracted from skin biopsies of old donors (>64 years old) compared to young donors (<32 years old) (Figures 5a, b, c), whereas *HDAC5* transcript levels were not significantly modulated (Supplementary Figure S1g). These results suggest that HDAC4 inhibition may mimic aging-related features in epidermal models. To further support this hypothesis, we analyzed several aging-associated gene expression (*SIRT1*, *IDH1*, *TP63 and CDKN1A* encoding P21) in RHE treated or not with LMK235 (Idda et al. 2020; Muther et al. 2017; Rivetti Di Val Cervo et al. 2012). As expected, we observed a significant decrease in *SIRT1*, *IDH1* and *TP63* mRNA levels, as well as an increase in *CDKN1A* mRNA levels upon treatment (Figure 5d), demonstrating that LMK235 can trigger the phenotype of an aged epidermis. Moreover, RNA-seq analysis revealed a significative down-regulation of *DNMT1*, *SERPINA3* and *TP53,* three other markers known to be repressed with aging (Choi et al. 2025; Ciccarone et al. 2016; Feng et al. 2007). It also confirmed the down-regulation of *SIRT1* associated with a significant up-regulation of *CDKN1A* (Figure 5e). Taken together, these results suggest that reconstructed epidermis subjected to HDAC4/5 inhibition exhibit several hallmarks of skin aging.

**Figure 5.**
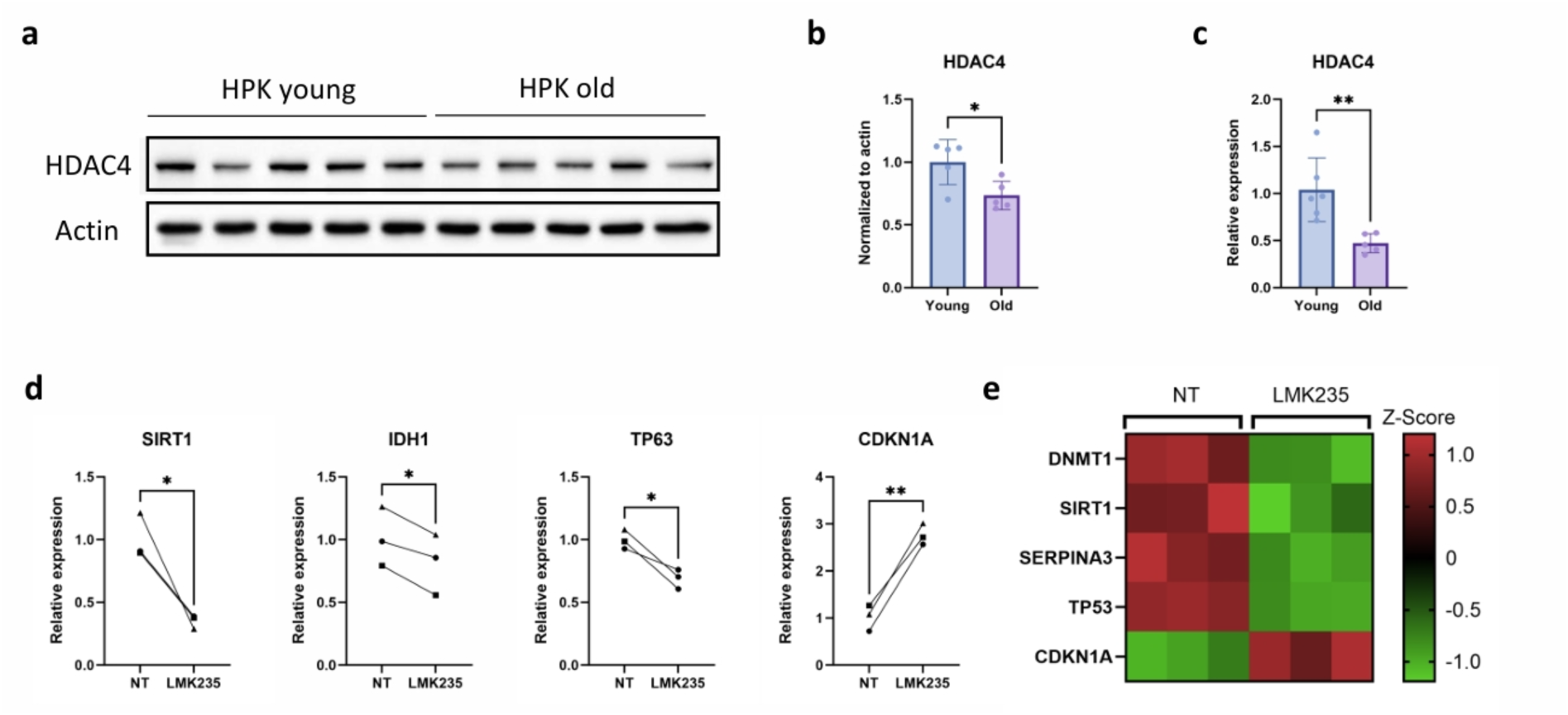
HDAC4 is downregulated during epidermal aging and its inhibition in reconstructed epidermis induces aging-associated hallmarks. (a) Western-blot analysis of HDAC4 in human primary keratinocytes (HPK) extracted from skin biopsies of young (<30 years old) and old donors (>64 years old). Actin was used as a loading control (N=5). (b) Quantification of HDAC4 protein levels in young and old HPK. Data were normalized to actin and expressed relative to the young HPK condition (* P<0.05, N=5, Student’s t-test). (c) Relative mRNA expression for *HDAC4* in young HPK (<32 years old) and old HPK (>64 years old). Data were normalized to housekeeping genes and expressed relative to the young HPK condition (** P<0.01, N=6 for young HPK and N=5 for old HPK, Student’s t-test). (d) Relative mRNA expression for *SIRT1*, *IDH1*, *TP63* and *CDKN1A* in reconstructed human epidermis treated or not with 1 µM LMK235. Data were normalized to housekeeping genes and expressed relative to the untreated condition (* P<0.05, ** P<0.01, N=3, Student’s t-test). (e) Heatmap representing the Z-score RNA-seq expression of *DNMT1*, *SIRT1*, *SERPINA3*, *TP53* and *CDKN1A*, 5 aging-associated genes, in reconstructed human epidermis treated or not with 1 µM LMK235. Normalized counts were log2-transformed and Z-scores were calculated for each gene across all samples using the formula Z=(x-mean)/SD where SD represents the sample standard deviation across samples. Each column represents a biological replicate (N=3) and each row represents a gene. Color intensity is proportional to the Z-score of each gene.

## DISCUSSION

In this study, we investigated the role of histone deacetylase HDAC4/5 in epidermal homeostasis and the underlying epigenetic mechanisms, in 3D models of reconstructed human epidermis and reconstructed full thickness skin. We showed that HDAC4/5 are essential to maintain proper keratinocyte proliferation and differentiation notably through their role in reshaping epigenetic landscape signature of these two cellular states. Additionally, our study led to the discovery of a potential role for these enzymes in controlling skin aging. Since relatively few relevant *in vitro* models mimicking intrinsic aging are available to date, a new model based on HDAC inactivation will certainly be of great interest for both basic and applied research (Briganti et al. 2026).

When considering this study from a broader perspective, it is interesting to note that transcriptional analysis by RNA-seq revealed that increased histone H3 acetylation by LMK235 treatment induced a drastic effect on global transcriptional profiles. Although we expected to observe predominantly up-regulated genes, since histone acetylation is generally associated with an open-chromatin state, we surprisingly identified a similar proportion of up and down-regulated genes. Similar observations have previously been reported in a human leukemia cell line, where the establishment of a more permissive chromatin state through increased histone acetylation following pan-HDAC inhibition did not result in global transcriptional activation (Slaughter et al. 2021). Indeed, a recent study highlighted the contribution of additional regulatory mechanisms involved in the control of gene expression changes, including the loss of active epigenetic marks (H3K27ac and H3K4me3), the enrichment of transcriptional regulators targeted by HDACs (Myc and YY1) and the formation or reinforcement of repressive chromatin loop at downregulated TSS (Paldi et al. 2026). Accordingly, we also showed that genes involved in lipid biosynthetic pathways were hyperacetylated in differentiated cells upon LMK235 treatment, despite their decreased transcript levels, as observed for *ELOVL4, ACER1* and *ALOX12B*. This suggests that the strong repression of lipid synthesis was not directly driven by epigenetic regulation but rather reflects a broader adaptative response to a metabolite-deprived condition. Indeed, in cancer cells, Basseville et al. reported that HDAC inhibitor treatment induces acetyl-CoA depletion (Basseville et al. 2022), an early precursor of *de novo* lipid synthesis (Gao et al. 2016). Herewith, LMK235 treatment might increase retention of acetyl groups on nuclear histones, thereby reducing the pool of free acetate and consequently acetyl-CoA availability.

Given the role of epidermal stem cells in skin renewal and considering the reduced proliferative capacity observed following HDAC4/5 inhibition, these enzymes might be involved in the maintenance of a progenitor-like cell state. Notably, our RNA-seq analysis revealed the down-regulation of several genes associated with the basal/progenitor compartment such as *FOXM1*, *KRT15*, *BMI1* and *NOTCH1*. *FOXM1* is a transcription factor able to maintain stem cell renewal through regulation of pluripotency-associated genes. It had been shown to target *KRT15*, a progenitor stem-like population marker and both genes are upregulated in basal cell carcinomas (Bose et al. 2013; Redmond et al. 2025). In addition, *NOTCH1* and *BMI1* contribute to the maintenance of stemness and interestingly are decreased during aging in human keratinocytes (Cordisco et al. 2010; Palazzo et al. 2015). These results support the hypothesis of (Roca et al. 2019) regarding HDAC inhibitors capacity to shift cancer cells from a stem like resistant state to a more differentiated and sensible phenotype thereby highlighting their therapeutical potential.

Whilst we have combined pharmacological inhibition with gene silencing of HDAC4, the silencing of HDAC5 could provide further insights in the specific role of each of these enzymes. Moreover, the specificity of LMK235 toward HDAC4/5 remains to be fully established and does not exclude off-target effects on other HDACs, particularly other class IIa HDACs as the hydrophobic dimethyl-substituted phenyl ring of LMK235 shows greater compatibility with this HDAC subfamily than with class I HDACs (Di Giorgio et al. 2015). Also, we here focused exclusively on histone H3 acetylation. Although this epigenetic mark is a well-established target of HDAC4/5 (Di Giorgio et al. 2020; Wang et al. 2026), this is not enough to capture the full complexity of epigenetic modifications changes involved in maintaining epidermal homeostasis. As illustrated by recent single-nuclei proteomic data in human keratinocytes, providing a global overview of histone variants specific to keratinocyte differentiation, additional changes in histone H4 acetylation also take place. As these modifications appear to be even more specific to the differentiated state than acH3, it highlights the importance of considering other histone acetylation marks in future studies (Kanchustambham et al. 2026). Moreover, even if the H3K27ac ChIP-seq analysis provides valuable insights into the impact of HDAC4/5 inhibition on H3K27ac-enriched genomic regions, profiling HDAC4/5 occupancy would be valuable to identify genomic regions directly targeted by these enzymes.

Collectively, our results show that HDAC4/5 are essential for maintaining epidermal homeostasis notably through their role in keratinocytes proliferation and epidermal barrier function. This study provides new insights into the role of epigenetic regulators in the epidermis and suggest that the dysregulation of the epigenetic landscape, in particular histone acetylation, may be one of the hallmarks and mechanisms of cellular aging.

## METHODS

### Cell culture

Human primary keratinocytes were isolated in-house from skin biopsies that were obtained from the DermoBioTec tissue bank at Lyon or from Biopredic International with the informed consent of adult donors (non-pathological tissues from abdomen or breast), in accordance with French ethical guidelines (French Bioethics law of 2021). Donor specifications are indicated in Supplementary Table S1. N/TERT1 keratinocytes cell line was originally generated by Jim Rheinwald (Dickson et al. 2000) and was kindly provided to us by Romain Debret (Lyon, France). HPK and N/TERT1 were cultured in supplemented KBM Gold medium (#00192152 et #00192151, Lonza) at 37 °C and 5% CO2. The culture medium was renewed three times a week. HPK and N/TERT1 were maintained at no more than 4 and 37 passages, respectively. The differentiation of HPK and N/TERT1 was induced by leaving the cells at confluency in supplemented KBM Gold medium.

### Reconstructed Human Epidermis

Reconstructed human epidermis (RHE) was generated on a translucent polycarbonate membrane. Briefly, adult human keratinocytes seeded at 500 000 cells/cm² were cultured for 2 days in Epilife Calcium-free and phenol red-free medium (MEPICFPRF500, Gibco) supplemented with 1.5 mM Ca^2+^ at 37 °C and 5 % CO2. Thereafter, the cultures were lifted to the air-liquid interface (D0) and maintained for 12 days in same basal medium supplemented with 1.5 mM Ca^2+^, 92 µg/mL L-Ascorbic acid 2-phosphate and 10 ng/mL KGF with medium changes performed every two days. RHE were treated with 1 µM LMK235 from day 4 to day 12, with inhibitor added at each medium change.

### Full thickness reconstructed human skin

To generate full thickness reconstructed human skin (RHS), human dermis equivalents were first produced by seeding human primary fibroblasts (D0) into an acellular matrix composed of collagen, chitosan and glycosaminoglycans (Mimedisc) and cultured in DMEM/F12 supplemented with 10 % Fetal Bovine Serum (FBS) and antibiotics, with medium renewal every two days. After 28 days, human primary keratinocytes were seeded onto the surface of the dermal equivalents and cultured for 9 days. Then, cultures were lifted to the air-liquid interface and maintained in DMEM/HAM F12 supplemented with 0.8 % FBS, 50 µg/mL ascorbic acid, hydrocortisone, insulin and antibiotics. The culture medium is renewed every two days. RHS were treated with 1 µM LMK235 from day 43 to day 49, with inhibitor added at each medium change.

### Chemical compounds

The HDAC4 inhibitor, LMK235 (AB-M4986, Ab Moles) is solubilized in DMSO to a final concentration of 91.7 mM. During the experiments, the inhibitor is used at a concentration of 1 µM.

### HDAC4 silencing

N/TERT1 were transduced with lentiviral vectors from scrambled Human shRNA plasmid kit (TR30021, Origen) and HDAC4 Human shRNA plasmid kit (#TR312493-ORIGEN). Lentiviral vector particles were produced by the AniRA lentivectors production facility from the CELPHEDIA Infrastructure and SFR Biosciences (UAR3444/CNRS, US8/Inserm, ENS de Lyon, UCBL). N/TERT1 cells were transduced 24h after seeding them at low density (5 000c/cm²) with lentiviral particles (MOI 200) containing a vector with a non-effective shRNA sequence (shRNA-SCR) or a vector with a shRNA targeting HDAC4 (sh-HDAC4) for 24h. Transduced cells were cultured under puromycin selection at 5 µg/mL for 4 days and then selected cells were amplified before analysis.

HPK were transfected with 25 nM ON-TARGETplus non-targeting siRNA (#D-001810-10-20-Dharmacon) or HDAC4-targeting siRNA (#J-003497-07-Dharmacon) using DharmaFECT 1 Transfection Reagent (#T-2001-02-Dharmacon) from Horizon Discovery, according to the supplier’s instructions.

### Cell viability assay

Cell viability assays in N/TERT1 cells treated or not with 1 µM LMK235 were performed using the kit AlamarBlue assay (DAL1100, Invitrogen) according to supplier’s instructions. Fluorescence was measured with the Spark microplate reader (Tecan) using an excitation wavelength of 560 nm and an emission wavelength of 590 nm. Cell viability was calculated as a percentage relative to control cells which were set to 100% of viability.

### Cell proliferation Assay

Cell proliferation assays in N/TERT1 cells treated or not with 1 µM LMK235 were performed using the kit CyQUANT Cell proliferation Assays (C7026, Invitrogen) according to supplier’s instructions. Fluorescence was measured with the Spark microplate reader (Tecan) using an excitation wavelength of 480 nm and an emission wavelength of 520 nm. Cell proliferation was quantified by comparing the obtained fluorescence values of each condition to a standard curve.

### HDAC activity Assay

HDAC activity assays in N/TERT1 cells treated or not with 1 µM LMK235 were performed using the kit Epigenase HDAC Activity/Inhibition Direct Assay (P-4034, EPIGENTEK) according to supplier’s instructions. 5 µg of protein were used for each reaction and the enzymatic reaction was carried out for 90 minutes. Absorbance was measured at 450 nm using the Spark microplate reader (Tecan).

### Skin tissue or RHE preparation

Skin tissues or RHE were fixed in 4 % paraformaldehyde (PFA) overnight at 4°C. Upon three washes, samples were dehydrated through successive ethanol baths of increasing concentrations (30 %, 50 % and 70 %). Tissue clearing and paraffin infiltration were subsequently performed using the Histo 5 Rapid Microwave HistoProcessor (MM France). The clearing step was carried out using the MW Unit with a tissue-specific program. Then, tissues were dried in the Wax module for 1’30 minutes using the “Vaporization” program, followed by paraffin impregnation using the “Impregnation” program. Paraffin embedding was performed thanks to the EC 350 embedding station (MM France). Tissue sections of 5 µm thickness were prepared using a Microm HM355S (MM France).

### Immunofluorescence straining

Following dewaxing and rehydration, tissue sections went through a pretreatment depending on the target antigen: permeabilization for 10 minutes at RT in a solution of 0.1 % Triton X-100 and 0.1 M glycine diluted in 1X PBS or antigen retrieval in boiling citrate buffer (1.8 mM citric acid monohydrate and 8.2 mM tri-sodium citrate dihydrate) for 20 minutes at exactly 98 °C. After three washes, sections were blocked for at least 1h at RT in blocking solution containing 5 % normal goat serum, 2 % BSA and 0.1 % Tween20 diluted in 1X PBS. Primary antibodies diluted in a solution of 2.5 % normal goat serum, 1 % BSA and 0.05 % Tween20 were incubated overnight at 4 °C. After three washes, secondary antibodies diluted in the same solution were incubated for 1h at RT in the dark. After three washes, nuclear staining and sections mounting were performed using Fluoroshield mounting medium containing DAPI (F6057, Sigma Aldrich). All primary and secondary antibodies used are listed in Supplementary Table S2. Immunofluorescence staining on cultured cells followed the same protocol, with 4 % PFA fixation for 10 min prior to staining and with the omission of the dewaxing and rehydration steps. Images were acquired using a motorized Nikon Ti-E inverted fluorescence microscope with the following objectives: Plan Fluor 10x Ph1 DL, Plan fluor ELWD 20x Ph1 ADM et S Plan Fluor EL WD 20x ph2 ADM. Image analysis and quantification were performed using Image J (version 1.54f) and QuPath (version 0.5.1).

### RNA extraction and real time PCR

Total RNA extraction from keratinocytes or RHE was performed using the Nucleospin RNA plus kit (740984, Macherey Nagel) according to manufacturer’s instructions. Extracted RNA were quantified with a NanoDrop ND-2000 spectrophotometer. Then, 500 ng of RNA was reverse-transcribed using the PrimeScript RT Master Mix (Perfect Real Time) (RR036A, Takara) following the supplier’s instructions, with the amplification performed with a PCR T100 Thermal cycler (Bio-Rad). Quantitative PCR (qPCR) was carried out using the TB Green Premix Ex Taq II (Tli RNaseH Plus) Bulk kit (RR820L, Takara) on an Azure Biosystem (Azure Cielo). Primer sequences are listed in Supplementary Table S3. Relative gene expression levels were calculated using the comparative Ct method (2^-ΔΔCt) and normalized to the expression of housekeeping genes. *PUM1* and *UBC* were used as housekeeping genes for the RHE experiments. *RPL13A* was used as the housekeeping gene for 2D models generated with untreated HPK and transduced N/TERT1. PUM1 and OAZ1 were used as housekeeping gene for LMK235-treated N/TERT1 monolayer cells.

### Protein Extraction and Western-blot

Cells proteins were extracted using RIPA buffer supplemented with protease inhibitors. After quantification with the kit Pierce BCA Protein Assay (23225, Thermo Scientific), 15 µg of proteins were loaded onto 10 or 15 % SDS-polyacrylamide gels and transferred onto PVDF Immobilon-P membrane (IPHV00010, Sigma Aldrich). Membranes were blocked for at least 1h30 under agitation at RT in a blocking solution (5 % milk, 0.1 % Tween20 diluted in 1X TBS) and then incubated overnight under agitation at 4 °C with primary antibodies diluted in the blocking solution. Upon three washes, membranes were incubated for 1h under agitation at RT with HRP-conjugated secondary antibodies diluted in blocking solution. After washing, protein bands were detected using SuperSignal West Pico PLUS substrate (34578, Thermo Scientific) and imaged with a Fusion Fx system (Vilber Lourmat). Primary and secondary antibodies used are listed in Supplementary Table S4.

### Lipid analysis

All lipid analysis were performed by QIMA. Briefly, 50 mg of full thickness reconstructed human skin, precisely weighted were used to perform liquid/liquid lipid extraction (following a protocol inspired from the Bligh & Dyer method). Cholesterol content was determined by liquid chromatography coupled to mass spectrometry (LC-MS). Fatty acids profiling was performed after derivatization by gas chromatography coupled to mass spectrometry (GS-MS). Ceramide analysis was conducted by liquid chromatography coupled to tandem mass spectrometry (LC-MS/MS) after solid-phase extraction. This method enabled semi-quantitative analysis of distinct ceramide subclasses (NS/AS and EOS).

### Functional analysis of human reconstructed epidermis

Trans-epidermal water loss of RHE was measured using a Tewitro TW24 device (CK ELECTRONIC) in an unventilated room. Measurements were recorded every second over a 15 min period. Trans-epidermal electrical resistance was measured using an EVOM Manual device (EV-MT-03-01, WPI) equipped with an STX4 electrode (EVM-EL-03-03-01, WPI). Measurements were performed by immersing one electrode into the insert and placing the second electrode outside the insert. All measurements were performed in triplicate for each sample.

### RNA sequencing

Total RNA collected from reconstructed human epidermis treated or not with 1 µM LMK235 were quantified using a Tape Station and the TapeStation Analysis Software 5.1. RNA-seq library preparation, sequencing and bioinformatics analysis were performed by Genewiz from Azenta Life Science. Briefly, mRNA was enriched using Poly-A selection, fragmented and reverse-transcribed into cDNA. After end repair, dA-tailing and adaptor ligation, libraries were amplified by PCR and purified. Sequencing was performed on an Illumina NovaSeq (paired-end, 150 bp, 30 million reads). Reads were trimmed to remove adapter sequences and low-quality bases using fastp v.0.23.2. Trimmed reads were mapped to the Homo sapiens hg38 reference genome using STAR aligner v.2.5.2b. Gene-level counts were generated using featureCounts (Subread package v.1.5.2). Only uniquely mapped reads overlapping exon regions were kept, and counts were summarized at the gene-id level based on the annotation file. Differential expression analysis was performed using DESeq2. The Wald test was used to generate p-values and log2 fold changes. Genes with an adjusted P<0.05 and |log2 fold change| >1 were considered differentially expressed.

### Transmission electron microscopy (TEM)

TEM was performed on reconstructed human epidermis treated or not with 1 µM LMK235. Tissues samples were fixed in 2% glutaraldehyde and 0.1 M cacodylate (pH 7.4, EM grade) at 4°C for several days. TEM samples preparation and images acquisition were performed at CIQLE platform (Centre d’Imagerie Quantitative Lyon Est, platform UCBL). Briefly, samples were washed three times for 1h at 4°C and contrasted with tea Oolong in 0.2 M cacodylate (pH 7.4, EM grade) for 1h at RT. Post-fixation was carried out using 1% aqueous osmium tetroxide (EMS) for 1h at RT. After three washes in 0.2 M sodium cacodylate followed by one wash in water, samples were dehydrated through a graded ethanol series and transferred to propylene oxide. Impregnation was performed with Epon A (75%), Epon B (25%) and DMP30 (1,7%). Inclusion was obtained by polymerization for 72h at 60 °C. Sections were made on a UC7 ultramicrotome (Leica Microsystems). Regions exhibiting alterations were first identified on semi-thin sections (1000 nm) stained with methylene blue Azur II. Ultrathin sections (approximately 70 nm) were then cut, mounted on 200 mesh copper grids coated with 1:1 000 poly-lysine, and stabilized for 24h at RT. Sections were subsequently contrasted with uranyl acetate and lead citrate, and examined with a JEOL 1400-JEM transmission electron microscope (Tokyo,Japan), operated at 100 kV, equipped with a Orius 1000 camera and Digital Micrograph software (version: 1.7).

### ChIP H3K27ac, library construction, Chip-seq and data analysis

N/TERT1 proliferative or differentiated cells treated or not with 1 µM LMK235 were fixed in 1% formaldehyde for 15 minutes at RT. Fixation was stopped by the addition of 0.125 M glycine (final). After washes in PBS-0.5% Igepal, fixed cell pellets were snap-frozen and sent to Active Motif Services (Carlsbad, CA) for ChIP-Seq processing. Active Motif performed chromatin preparation, immunoprecipitation, library construction, and sequencing. Briefly, chromatin was isolated using lysis buffer and sheared by sonication using a PIXUL® Multi-Sample Sonicator (53130, Active Motif) to obtain DNA fragments with a size of 200–1000 bp. To determine chromatin yield, an aliquot of sheared chromatin was reverse crosslinked at 65°C, treated with RNase and proteinase K, and submitted to DNA purification using SPRI beads (Beckman Coulter). DNA concentrations were measured using a Qubit Fluorometer (Thermo Fisher), and total chromatin yield was extrapolated based on the original chromatin volume. For immunoprecipitation, chromatin samples were precleared with protein G agarose beads (Invitrogen). ChIP was performed using 4 µg of anti-H3K27ac antibody (39133, Active motif) per 13 µg of chromatin for each reaction. After washing, immune complexes were eluted from the beads using SDS buffer, RNase and proteinase K, and reverse crosslinking was performed by overnight incubation at 65°C. ChIP DNA was then purified using phenol-chloroform extraction and ethanol precipitation. ChIP DNA libraries were prepared using the NEB DNA Library Prep Kit, following the manufacturer’s instructions. Illumina sequencing libraries (a custom type, using the same paired read adapter oligonucleotides described by (Bentley et al. 2008)) were prepared from the ChIP and Input DNAs on an automated system (Apollo 342, Wafergen Biosystems/Takara). After final PCR amplification, libraries were quantified and sequenced on Illumina’s NovaSeq (paired-end, 50 bp, 30 million reads). Then, data processing and analysis were performed by Altrabio (Lyon, France). Sequencing reads were trimmed and mapped on the “primary assembly” Homo sapiens genome sequence (release GRCh38.p14). Filtering steps removed reads mapping to ENCODE blacklist regions, duplicate reads, unmapped reads, non-primary alignment, multi-mapping reads, reads with more than 4 mismatches and discordant read pairs. Peak calling was performed using Omnipeak with Input sample as control applying a false discovery rate threshold of 0,05. Normalization factors were calculated for each sample using a combination of DESeq2 Size Factor and ChIPseqSpikeInFree estimation tools.

### Statistical analysis

Data are expressed as mean ± SD. Depending if our data set follow a normal distribution (Shapiro-Wilk test), statistical significance was calculated by Student’s T-test or Mann-Whitney and two-way analysis of variance (ANOVA2) using Prism software (version 10.1.1, GraphPad Software, San Diego, CA, USA). Mean differences were considered statistically significant when *P* < 0.05 (* *P* < 0.05, ** *P* < 0.01, *** *P* < 0.001, **** *P* < 0.0001).

## ACKNOWLEDGMENTS

We thank Charlène Prier for support in the development of the full thickness reconstructed human skin and Audrey Pierrot for assistance with staining procedures and visualization. We acknowledge the contribution of CIQLE platform (Centre d’Imagerie Quantitative Lyon Est, platform UCBL) for the preparation of transmission electron microscopy samples and image acquisition. We acknowledge the contribution of AniRA lentivectors production facility from the CELPHEDIA Infrastructure and SFR Biosciences (UAR3444/CNRS, US8/Inserm, ENS de Lyon, UCBL), especially Gisèle Froment, Aurélie Thibaut and Caroline Costa. We thank the PrImaTiss platform (Tissue preparation and imaging platform, LBTI) and more specifically Sandra Ferraro, for providing practical training, access to the platform’s equipment, validated protocols, and technical support. We also thank BioMolTools platform (Molecular Biology platform, LBTI) for providing practical training, access to the platform’s equipment, validated protocols and valuable advice. We thank our valuable trainee Jeanne Grangy for her contribution to the development and optimization of several protocols. We thank Pr. Violaine Sée and Lisa Martin for the critical reading of the manuscript.

## AUTHOR CONTRIBUTIONS

Conceptualization: JL, VA, NP; Data Curation: CN; Formal Analysis: CN; Funding Acquisition: JL, NP, VA; Investigation: CN, SD, SC; Methodology: CN, SD, SC, NP, VA, JL; Project Administration: CN, NP, VA, JL; Resources: SD, SC, NP; Supervision: NP, VA, JL; Validation: CN, NP, VA, JL; Visualization: CN; Writing - Original Draft Preparation: CN, JL; Writing - Review and Editing: CN, SD, SC, NP, VA, JL

