## supplementary data for "HDAC4/5 regulate epidermal barrier function by modulating the epigenetic landscape of human keratinocytes"

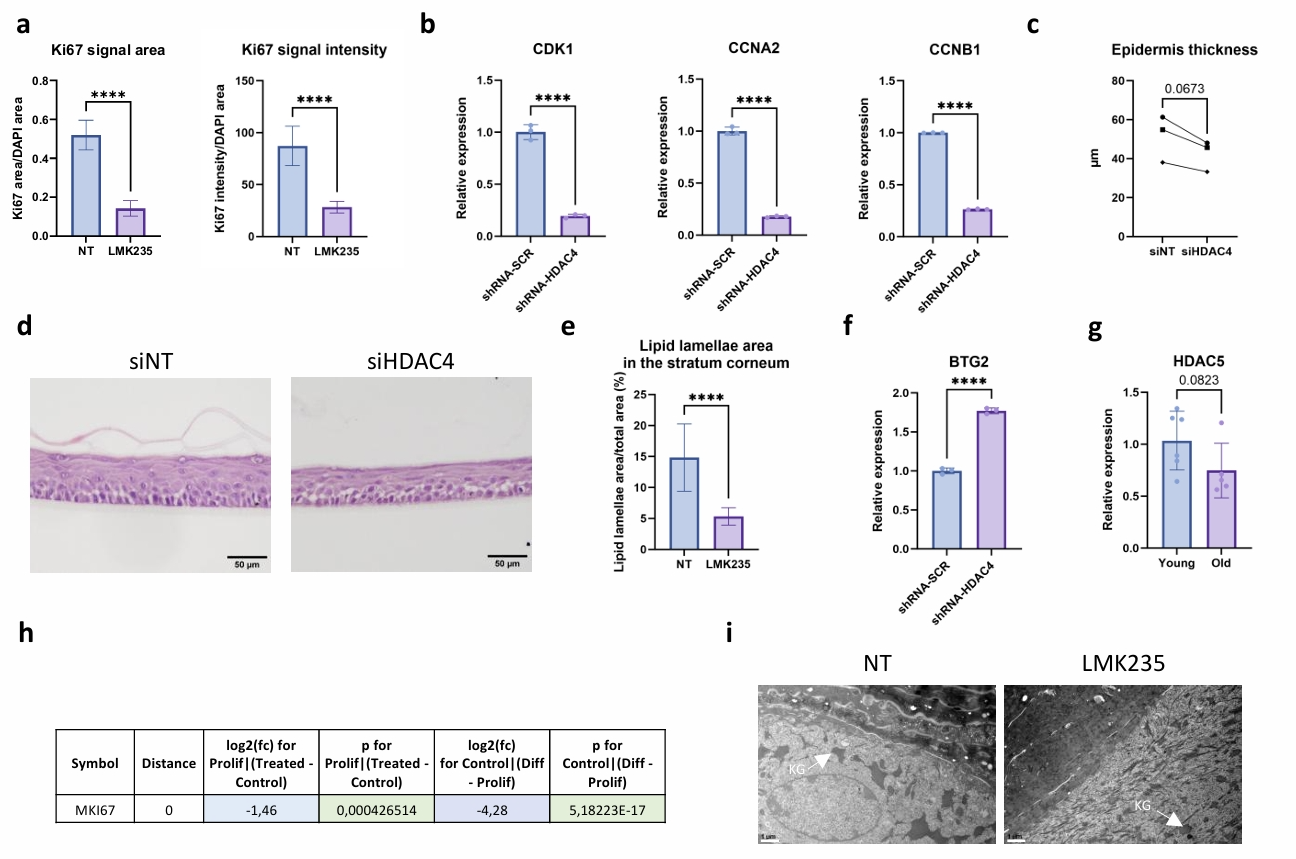
**Supplementary Figure S1:** (a) Quantification of the Ki67 signal area and signal intensity in N/TERT1 monolayer cells treated or not with 1 µM LMK235 for 24h (**** P<0.0001, N=1, n=15, Student’s t-test). (b) Relative mRNA expression for *CDK1*, *CCNA2* and *CCNB1* in N/TERT1 monolayer cells transduced with shRNA-SCR or shRNA-HDAC4. Data were normalized to housekeeping genes and expressed relative to the untreated condition (**** P<0.0001, n=3, Student’s t-test). (c) Representative H&E images of reconstructed human epidermis generated from HPK transfected with siNT and siHDAC4. Bars= 50 µM. (d) Quantification of epidermis thickness in reconstructed human epidermis generated from HPK transfected with siNT and siHDAC4 (N=3, Student’s t-test). (e) Quantification of the lipid lamellae area in the stratum corneum of reconstructed human epidermis treated or not with 1 µM LMK235 (**** P<0.0001, N=1, n=45, Student’s t-test). (f) Relative mRNA expression for *BTG2* in N/TERT1 monolayer cells transduced with shRNA-SCR or shRNA-HDAC4. Data were normalized to housekeeping genes and expressed relative to the untreated condition (**** P<0.0001, n=3, Student’s t-test). (g) Relative mRNA expression for *HDAC5* in young HPK (<32 years old) and old HPK (>64 years old). Data were normalized to housekeeping genes and expressed relative to the young HPK condition (N=6 for young HPK and N=5 for old HPK, Mann-Whitney test). (h) Table showing the differential H3K27ac peak analysis at the transcription start site (TSS; distance =0) of the *MKI67* gene. The table shows the Log2 Fold Change (log2(fc)) and p-value (p) for the comparison LMK235 treated and untreated proliferative NTERT1 cells (Prolif|(Treated-Control)) and between untreated differentiated and proliferative N/TERT1 cells (Control|(Diff-Prolif)). Negative log2fc values indicate hypoacetylation of H3K27 whereas positive log2fc values indicate hyperacetylation in the first condition relative to the second of each comparison. (i) Transmission Electron Microscopy (TEM) representative images of the *stratum granulosum* of reconstructed human epidermis treated or not with 1 µM LMK235. KG: keratohyalin granules. Bars= 1 µm.

***Supplementary table S1. Donor characteristics of the human primary keratinocytes used.***

| **# donor** | **Anatomical location** | **Age** | **Sex** | **Phototype** |
| --- | --- | --- | --- | --- |
| HPK20060 | Abdomen | 30 | F | V |
| HPK11197 | Breast | 26 | F | - |
| HPK22045 | Abdomen | 32 | F | II/III |
| HPK23094 | Abdomen | 30 | F | II/III |
| HPKS2L060 | Abdomen | 29 | F | Caucasian |
| HPKSL069 | Abdomen | 27 | F | Caucasian |
| HPK24041 | Abdomen | 30 | F | Caucasian |
| HPK24039 | Abdomen | 25 | H | II/III |
| KA20043 | Abdomen | 24 | F | VI |
| HPK24008 | Abdomen | 70 | F | II/III |
| HPK20055 | Breast | 64 | F | II/III |
| HPK23096 | Abdomen | 67 | F | II/III |
| HPKS2L073 | Abdomen | 65 | F | Caucasian |
| KS11162 | Breast | 68 | F | - |
| HPK240 | Foreskin | Child | H | - |
| HPK254 | Foreskin | Child | H | - |
| HPK21090 | Foreskin | Child | H | - |
| HPK21091 | Foreskin | Child | H | - |
| HPK-BASF | Abdomen | 37 | F | - |
| HPF-BASF | Breast | 18 | F | - |

***Supplementary table S2. List of antibodies used for immunofluorescence staining.***

| **Primary antibody** | | | |
| --- | --- | --- | --- |
| **Target** | **Origin** | **Antibody reference** | **Dilution used** |
| acH3 | Rabbit | 06-599 (MILLIPORE) | 1 : 300 |
| PCNA | Mouse | SC-56 (SANTA CRUZ) | 1 : 500 |
| Ki67 | Mouse | MAB4190 (SIGMA-ALDRICH) | 1 : 450 |
| loricrin | Rabbit | Ab85679 (ABCAM) | 1 : 500 |
| involucrin | Mouse | SC-21748 (SANTA CRUZ) | 1 : 500 |
| K14 | Rabbit | Ab181595 (ABCAM) | 1 : 500 |
| **Secondary antibody** | | | |
| **Target** | **Origin** | **Antibody reference** | **Dilution used** |
| Anti-rabbit IgG 546nm | Goat | A11035 (INVITROGEN) | 1 : 900 |
| Anti-mouse IgG2a 546nm | Goat | A21133 (INVITROGEN) | 1 : 900 |
| Anti-mouse IgG1 546nm | Goat | A21123 (INVITROGEN) | 1 : 900 |

***Supplementary table S3. List of primer sequences used for RT-qPCR analysis.***

| **Target** | **Forward sequence** | **Reverse sequence** |
| --- | --- | --- |
| *HDAC4* | GAGAGACTCACCCTTCCCG | CCGGTCTGCACCAACCAAG |
| *PCNA* | CCTTGAGTGCCTCCAACACC | GACTTTCCTCCTTCCCGCC |
| *MKI67* | CTTTGGGTGCGACTTGACG | GTCGACCCCGCTCCTTTT |
| *IVL* | TCCTCCAGTCAATACCCATCAG | GCAGTCATGTGCTTTTCCTCTTG |
| *LOR* | CAGAACTAGATGCAGCCGGAGA | TCATGATGCTACCCGAGGTTTG |
| *CCNA2* | GAAGACGAGACGGGTTGCA | AGGAGGAACGGTGACATGCT |
| *CCNB1* | ATAAGGCGAAGATCAACATGGC | TTTGTTACCAATGTCCCCAAGAG |
| *CCND1* | CCTCTAAGATGAAGGAGACCA | AAATGAACTTCACATCTGTGGC |
| *CDK1* | AAACTACAGGTCAAGTGGTAGCC | TCCTGCATAAGCACATCCTGA |
| *SIRT1* | TGCTGGCCTAATAGAGTGGCA | CTCAGCGCCATGGAAAATGT |
| *IDH1* | TTGGCTGCTTGCATTAAAGGTT | GTTTGGCCTGAGCTAGTTTGA |
| *CDKN1A* | AGCTGCCGAAGTCAGTTCCTT | GTTCTGACATGGCGCCTCCT |
| *TP63* | AAGAAAGGACAGCAGCATTGAT | GGGACTGGTGGACGAGGAG |
| *ELOVL4* | GAGCCGGGTAGTGTCCTAAAC | CACACGCTTATCTGCGATGG |
| *CERS2* | GGTAGAGCGTTGGTTCCGTC | GGCAATGAAGGCAATCAGGTAA |
| *CERS3* | CCATCCAGTAGCTTCGCCTC | TCAGAGAGCAGCTTCCAACG |
| *ASAH1* | ATTGGCCCCAGCCTACTTTAT | CCCTGCTTAGCATCGAGTTCAT |
| *BTG2* | ACCTGCAAGAACCAAGTGCT | AGGTATGTGGTGGCCTGTTG |
| *OAZ1* | GGATCCTCAATAGCCACTGC | TACAGCAGTGGAGGGAGACC |
| *PUM1* | CGGTCGTCCTGAGGATAAAA | CGTACGTGAGGCGTGAGTAA |
| *RPL13A* | CTCAAGGTCGTGCGTCTGAA | TGGCTGTCACTGCCTGGTACT |
| *UBC* | CACGTCAGACGAAGGGCGCAG | CGCCTGTTCCGCTCTCTGCTGGAAA |

***Supplementary table S4. List of antibodies used for Western-blot.***

| **Primary antibody** | | | |
| --- | --- | --- | --- |
| **Target** | **Origin** | **Antibody reference** | **Dilution used** |
| actin | Rabbit | Ab8227 (ABCAM) | 1 : 5000 |
| acH3 | Rabbit | 06-599 (MILLIPORE) | 1 : 500 |
| HDAC4 | Rabbit | GTX110231 (GENTEX) | 1 : 1000 |
| **Secondary antibody** | | | |
| **Target** | **Origin** | **Antibody reference** | **Dilution used** |
| HRP IgG anti-Rabbit | Goat | Ab6721 (ABCAM) | 1 : 10000 |
